# Self-enhancement and devaluation of others during belief formation are linked to narcissistic traits

**DOI:** 10.64898/2026.10.21.760328

**Authors:** Merle Matthiesen, Annalina V. Mayer, Alexander Schröder, Laura Rosenbusch, Jan Philipp Klein, Ines Wilhelm-Groch, David S Stolz, Frieder M Paulus, Laura Müller-Pinzler, Sören Krach, Nora Czekalla

**Affiliations:** Social Neuroscience Lab, Department of Psychiatry, Psychotherapy and Psychosomatics, University of Lübeck, Lübeck, Germany; Center of Brain, Behavior and Metabolism (CBBM), University of Lübeck, Lübeck, Germany; Department of Psychology, Julius-Maximilians-Universität, Würzburg, Germany; Department of Psychiatry, Psychotherapy and Psychosomatics, University of Lübeck, Lübeck, Germany

**Author notes:** Corresponding authors: Nora Czekalla, Sören Krach.

**Keywords:** narcissistic personality, belief-updating, devaluation, social learning, positivity bias, self-enhancement

## Abstract

**Background:** Narcissistic personality traits are associated with the tendency to maintain a grandiose self-image through self-enhancement and devaluation of others. Using computational modeling, we investigated how narcissistic traits and the experience of pride are linked to how individuals form beliefs about themselves and others.

**Methods:** Healthy individuals and individuals with psychiatric diagnoses varying in narcissistic traits (N = 97) completed a learning task in which self-serving biases emerged implicitly through trial-by-trial belief updating in response to manipulated feedback. Computational modeling quantified how participants updated beliefs about their own and another person’s abilities in response to better- and worse-than-expected performance feedback.

**Results:** Higher narcissistic traits were associated with reduced learning from worse-than-expected self-related feedback, consistent with self-enhancement as a mechanism for maintaining a grandiose self-image. Individuals with higher narcissistic traits also reported greater pride following better-than-expected self-related feedback, which mediated the association between narcissistic traits and positive self-related learning. Furthermore, higher narcissistic traits were, mediated via pride, linked to increased learning from worse-than-expected feedback about the other person, suggesting that devaluing others relative to oneself may serve as a secondary pathway for maintaining a grandiose self-image.

**Discussion:** These findings identify computational and affective mechanisms through which narcissistic traits are linked to self- and other-related belief formation. Reduced integration of negative self-related feedback together with enhanced pride following positive feedback may support grandiose self-beliefs, while enhanced learning from negative information about others suggests that individuals with higher narcissistic traits implicitly construct themselves as superior through both self-enhancement and devaluation of others during interpersonal belief formation.

## Introduction

Narcissism is characterized by the pursuit and maintenance of a grandiose self-image^1^. This goal can be reached via different routes: first, information can be processed in a self-enhancing way^2,3^, strengthening the belief that one is better than the average person^4^. Second, a grandiose self-image can be maintained through the devaluation of others relative to oneself^5^. The aim of this study was to examine the role of narcissistic personality traits in how individuals process feedback and thereby form new beliefs about themselves and others. We used a learning paradigm, the Learning Of Own Performance (LOOP) task^6–8^, in which participants update self- and other-related beliefs through trial-by-trial performance feedback.

Narcissism is a heterogeneous and multidimensional construct that subsumes a combination of distinct personality traits^9^. Fundamental to narcissism is a biased self-perception accompanied by a strong urge to present oneself favorably to the outside world. This can be achieved by presenting oneself in a grandiose manner, which goes hand in hand with an excessive need for admiration and self-idealization, as well as by belittling or devaluing others and thereby elevating oneself^10^. These traits exist along a continuum, ranging from those within the healthy personality spectrum to subclinical personality accentuations, and ultimately to manifestations of narcissistic personality disorder^11^. At the maladaptive end of this continuum, narcissistic traits are associated with impaired psychosocial functioning and increased vulnerability to psychopathology, including depression and interpersonal difficulties^10^.

Generally, the motive of self-enhancement also plays a central role in the context of self-related information processing and the formation of self-beliefs^12,13^. People develop beliefs about themselves over time by interacting with others, receiving and interpreting social feedback^14^. They generate predictions based on these interactions and seek confirmation of their self-image^15^. When there is a discrepancy between self-beliefs and external information, meaning that feedback is better or worse than expected, it prompts an effort to minimize this inconsistency by adapting the belief^16^. Here, new information can be integrated in a biased way^8,17^ so that positive and negative information is weighted and integrated differently into the self-concept^6,18–21^. Some studies report an updating bias towards desirable self-related information^18,20,22^, others find a bias towards negative information^6,8,19^. These contrasting findings may be explained by differences in learning context, with biases toward desirable information emerging for stable self-related characteristics^18,20^ and biases toward negative information arising when feedback depends on trial-by-trial performance and effort^6,8^.

Regardless of the learning context, people tend to interpret new information through the lens of their prior beliefs^6,16,23^. Since situation-specific predictions are continuously generated on the basis of more global internal models, prior beliefs play a central role in processing new information^14^. Consequently, individual differences in these underlying beliefs may contribute to variability in belief updating, making it important to examine how personality traits are associated with self- and other-related belief formation. Central to the present study is whether narcissism as a personality trait is associated with how self- and other-related beliefs are formed. Previous studies on the association between narcissism and feedback processing demonstrated that narcissism is related to greater sensitivity to interpersonal feedback^1^. When receiving negative feedback, individuals with high levels of narcissism are particularly likely to respond by denying its legitimacy and minimizing the competence of the rater^24^. This means that when the self is threatened by negative feedback, antagonistic self-protection activates a devaluation of other people^1,25^. Conversely, in response to positive feedback, individuals with high levels of narcissism tend to view the feedback as legitimate and the rater as competent and likable^24^. Desired feedback is then used as confirmation of one’s self-perception as being unique and grandiose^1^. Thus, with elevated narcissistic personality traits, self-related information processing is biased towards desirable feedback, which serves to maintain and reinforce positive self-assessments and to protect oneself from negative self-evaluations by devaluing others^24^.

Apart from individual differences in prior beliefs and personality, it has been shown that emotional experiences play a central role in the processing of self-related feedback^7,26^. This relationship is particularly relevant in the context of narcissism because narcissistic self-views are strongly tied to external evaluation and are accompanied by pronounced emotional responses to both affirmation and threat^27,28^. Specifically, negative evaluations by others may threaten the grandiose self-image characteristic of narcissism, thereby eliciting feelings of shame that are often defensively masked by exaggerated expressions of pride^27^. Also, desired feedback is perceived more strongly and accompanied by a sense of pride, thereby potentially helping to boost the grandiose self-image^27,29^. It can therefore be assumed that narcissistic needs and motives influence emotional reactions to either threats or affirmations of a positive self-image.

The present study aimed to examine the association between narcissistic personality traits and the formation of new beliefs about oneself and others and whether affective states mediate this association. For this, a clinical and non-clinical sample with varying degrees of narcissistic personality traits was reanalyzed regarding this question^8^. The clinical sample was recruited from a day clinic specializing in interpersonal problems in chronic depression, a condition often accompanied by personality pathology.

We hypothesized that self-belief updates in response to better- and worse-than-expected performance feedback would be biased in a self-enhancing way with increasing narcissistic traits. Specifically, we expected individuals with stronger narcissistic traits to weigh better-than-expected feedback more and worse-than-expected feedback less strongly when updating beliefs about themselves. In contrast, given that the devaluation of others relative to oneself may function to maintain and enhance one’s own self-image, we expected that higher levels of narcissistic traits would be associated with stronger weighting of worse-than-expected feedback and weaker weighting of better-than-expected feedback when updating beliefs about others. Finally, we assumed that pride experienced during self-related belief updating mediates the relationship between narcissistic traits and information processing.

## Results

### Measuring belief formation

We used the Learning Of Own Performance (LOOP) task^6,7^ to characterize learning processes underlying self- and other-related belief formation. In this task, *n*=97 participants received manipulated feedback about their own or another person’s estimation performance (e.g., estimating the weight of animals or the height of buildings; see Figure 1A). Each trial began with a cue indicating the estimation category (e.g., weight or height) and the target agent (Self or Other), followed by a rating of expected relative performance for oneself or for the other person. After the estimation task, participants received manipulated feedback on their own (Self) or the other’s performance (Other). For each agent, one estimation category was paired with predominantly better-than-expected feedback (i.e., positive prediction errors; High Ability), the other category with predominantly worse-than-expected feedback (i.e., negative prediction errors; Low Ability) to induce adjustments in trial-by-trial performance expectations. This resulted in four feedback conditions defined by Agent (Self vs. Other) and Ability (High vs. Low; see Figure 1B).

**Figure 1.**
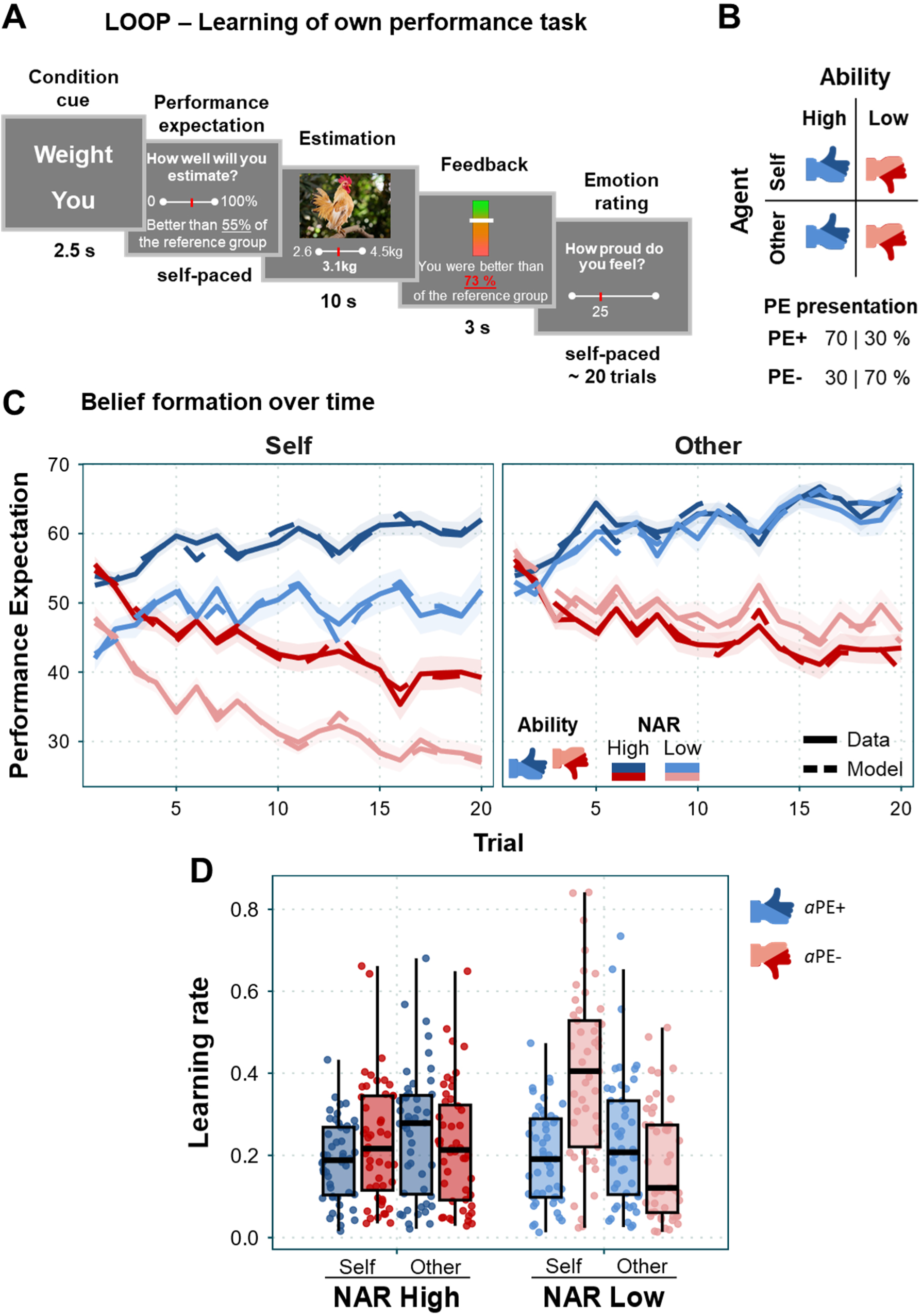
Trial sequence, experimental conditions, behavioral data, and learning rates. **A** Sequence of one self-related trial from left to right: At the beginning of each trial, a screen indicates the estimation category and agent of the upcoming trial. This is followed by a rating of the expected performance (performance expectation). Next, participants estimate specific attributes of the presented objects (estimation), after which they receive manipulated performance feedback (feedback). Approximately every 20 trials, participants are additionally asked to rate their current emotional state (e.g., pride). **B** The LOOP task consists of four distinct conditions. Low and High Ability conditions of alleged estimation performance for Self and Other (Agent condition) were presented in a pseudo-randomized order. Feedback with controlled prediction errors (PE): High Ability: 70 %, Low Ability: 30 % of the trials with planned positive PEs (the actual percentage of positive/ negative PEs could slightly differ, e.g., if the feedback had been out of range). **C** Performance expectations across 20 trials for all conditions (solid lines; shaded areas = ±1 SE). Dashed lines show predictions by the winning model (see Results), illustrating close correspondence with participants’ ratings. Narcissistic Admiration and Rivalry Questionnaire (NARQ) scores are median-split for display purposes; NAR = high vs. low narcissistic personality traits. **D** Model-derived learning rates for positive (*α*_PE+_) and negative prediction errors (*α*_PE-_) for self- and other-related learning. NARQ scores are median-split for display purposes.

In addition, participants rated their current emotional states (e.g., pride) four times during task performance. The participants’ level of narcissism was assessed with the Narcissistic Admiration and Rivalry Questionnaire (NARQ^1^, global scale).

### Higher performance expectations with higher narcissistic personality traits

To quantify how individuals adapt the performance expectation for themselves and the other person towards the presented feedback and how narcissistic personality traits alter this process, a model-free behavioral analysis was implemented first. Consistent with previous results, the Ability (High vs Low) × Agent (Self vs Other) × Trial × NARQ linear mixed model showed that participants adapted their performance expectation ratings according to the feedback (Ability × Trial interaction *t*(7363.9)=-20.81, *p*<.001), thus formed a new ability belief over time (see Fig. 1B). Moreover, participants evaluated their own performance increasingly more negatively compared to the other person’s performance (main effect Agent *t*(131.8)=-3.87, *p*<.001; Trial × Agent interaction *t*(7363.9)=-5.18, *p*<.001).

Individuals high in narcissistic personality traits had higher performance expectations compared to low narcissistic ones, specifically for themselves (Agent × NARQ interaction *t*(131.8)=2.25, *p*=.026). These higher self-related performance expectations increased over time (Agent × Trial × NARQ interaction *t*(7363.9)=3.26, *p*=.001), which can indicate greater self-belief updating after positive feedback or reduced self-belief updating after negative feedback (see Supplementary Table 1 for full results and Supplementary Table 2 for control analysis including group).

### Negativity bias in self-belief formation reduced with higher narcissistic personality traits

To quantify belief formation over time, performance expectation ratings were modeled using different variants of an adapted Rescorla-Wagner learning model (see Methods and previous work using the same task^6–8^). Model comparisons on the learning models indicated that a weighted Valence Model, which included separate learning rates for Self and Other, as well as positive (αPE+) and negative prediction errors (αPE-), best accounted for the expectation data (see Methods for the full model space; for model comparison and all PSIS-LOO scores, see Supplementary Table 3). The model was therefore selected for all subsequent analyses. As previous studies with the LOOP task have already shown, learning rates are higher for negative compared to positive prediction errors when updating self-beliefs but not for beliefs about the other person (PE-Valence × Agent interaction *t*(190)=-6.98, *p*<.001; post-hoc: Self αPE-vs. Self αPE+ *t*(96)=5.71, *p*<.001). This self-related negativity bias is reduced in individuals high in narcissistic personality traits (PE-Valence × Agent × NARQ interaction *t*(190)=3.36, *p*<.001; see Supplementary Table 4 for full results and Supplementary Table 5 for control analysis including group).

### Narcissistic traits predict self-enhancing and more negative learning about others via pride

We next tested whether narcissistic personality traits are associated with self- and other-related belief formation and whether the emotion of pride mediates this. To this end, we conducted mediation analyses using a path-analytic model (see Methods for details). The main model included narcissistic personality traits (NARQ) as a predictor, pride (mean pride ratings during learning) as a mediator, and the four learning rates for self- and other-related learning from positive vs. negative prediction errors as dependent variables (see Figure 2; for all correlations between variables, see Table 1).

**Figure 2.**
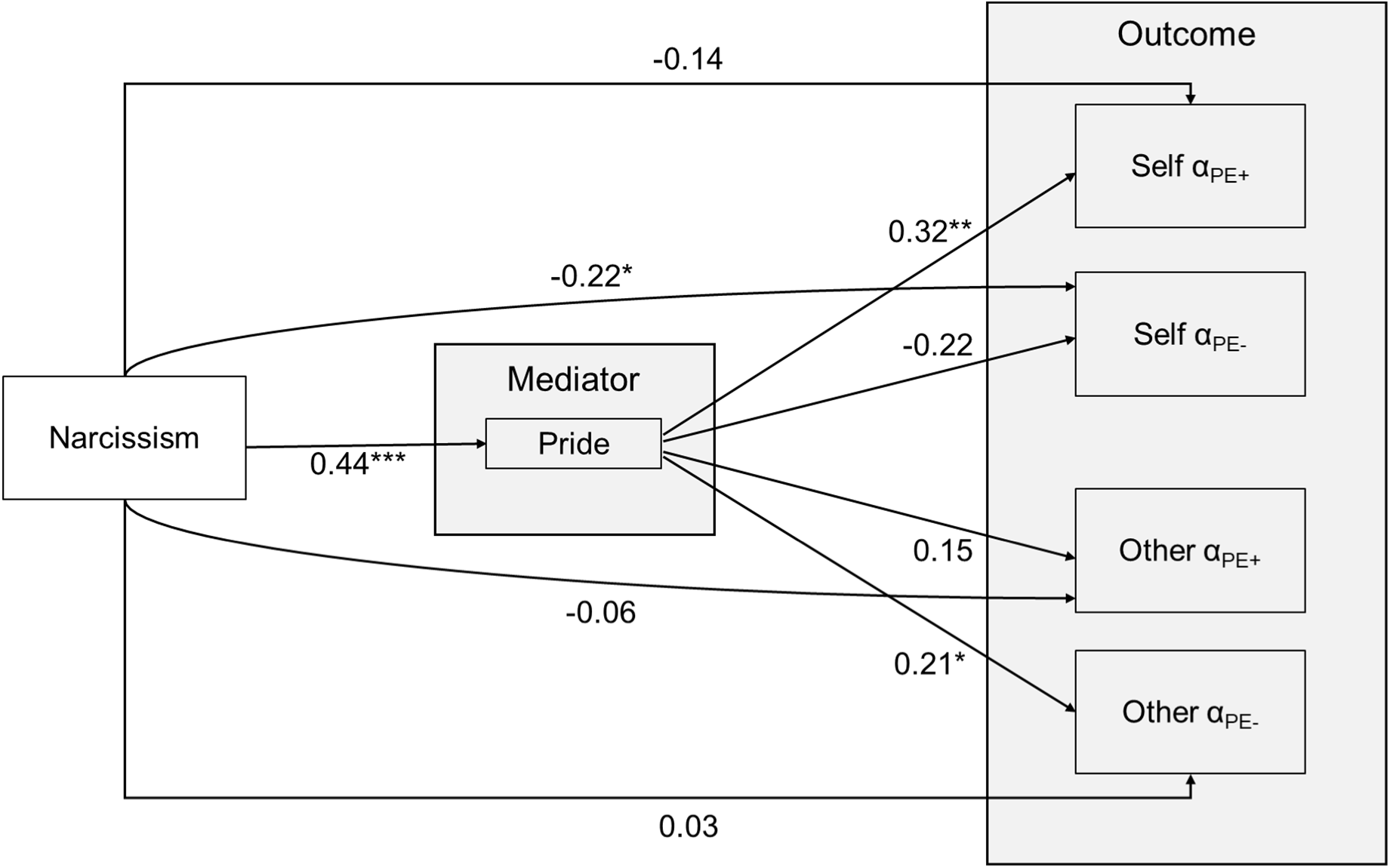
Path-analytic model illustrating the mediation of narcissistic traits (Narcissism) via pride (Mediator) on learning rates for positive (α_PE+_) and negative prediction errors (α_PE-_) in self-related (Self) and other-related (Other) learning. Arrows indicate direct effects; values represent standardized path coefficients. Indirect (mediation) effects and further statistics are reported in Table 2. * *p* < .05, ** *p* < .01, *** *p* < .001.

**Table 1.** Spearman correlations between variables.

| | NARQ | Pride | $\alpha_{PE+}$ Self | $\alpha_{PE-}$ Self | $\alpha_{PE+}$ Other | $\alpha_{PE-}$ Other |
| --- | --- | --- | --- | --- | --- | --- |
| NARQ | — |  |  |  |  |  |
| Pride | 0.436*** | — |  |  |  |  |
| $\alpha_{PE+}$ Self | 0.023 | 0.272** | — | | | |
| $\alpha_{PE-}$ Self | -0.293** | -0.308** | 0.205* | — | | |
| $\alpha_{PE+}$ Other | 0.063 | 0.154 | 0.315** | 0.220* | — | |
| $\alpha_{PE-}$ Other | 0.159 | 0.240* | 0.403*** | 0.198 | 0.688*** | — |

*Note.* Correlation table showing Spearman correlations for all variables, including all predictor, mediator, and dependent variables as used in the path-analytic mediation model. \* $p < .05$ , \*\* $p < .01$ , \*\*\* $p < .001$

**Table 2.** Path-analytic model.

| | *B* | *SE* | 95% CI | | *z* | *p* | $\beta$ |
| --- | --- | --- | --- | --- | --- | --- | --- |
|  |  |  | lower | upper |  |  |  |
| **Direct effects** |  |  |  |  |  |  |  |
| NARQ → Pride | 0.829 | 0.161 | 0.510 | 1.132 | 5.14 | < .001 | 0.44 |
| Pride → $\alpha_{PE+}$ Self | 0.034 | 0.012 | 0.011 | 0.055 | 2.89 | 0.004 | 0.32 |
| NARQ → $\alpha_{PE+}$ Self | -0.027 | 0.020 | -0.064 | 0.013 | -1.34 | 0.181 | -0.14 |
| Pride → $\alpha_{PE-}$ Self | -0.044 | 0.024 | -0.094 | 0.001 | -1.80 | 0.072 | -0.22 |
| **NARQ → $\alpha_{PE-}$ Self** | **-0.080** | **0.040** | **-0.158** | **-0.002** | **-2.00** | **0.045** | **-0.22** |
| Pride → $\alpha_{PE+}$ Other | 0.023 | 0.016 | -0.007 | 0.056 | 1.44 | 0.149 | 0.15 |
| NARQ → $\alpha_{PE+}$ Other | -0.018 | 0.032 | -0.078 | 0.050 | -0.57 | 0.571 | -0.06 |
| Pride → $\alpha_{PE-}$ Other | 0.029 | 0.014 | 0.004 | 0.058 | 2.16 | 0.031 | 0.21 |
| NARQ → $\alpha_{PE-}$ Other | 0.009 | 0.029 | -0.044 | 0.070 | 0.31 | 0.755 | 0.03 |
| **Indirect effects** |  |  |  |  |  |  |  |
| **NARQ → Pride → $\alpha_{PE+}$ Self** | **0.028** | **0.011** | **0.009** | **0.050** | **2.65** | **0.008** | **0.14** |
| NARQ → Pride → $\alpha_{PE-}$ Self | -0.036 | 0.022 | -0.084 | 0.001 | -1.64 | 0.102 | -0.10 |
| NARQ → Pride → $\alpha_{PE+}$ Other | 0.019 | 0.014 | -0.006 | 0.050 | 1.38 | 0.169 | 0.07 |
| **NARQ → Pride → $\alpha_{PE-}$ Other** | **0.024** | **0.012** | **0.003** | **0.048** | **2.09** | **0.037** | **0.09** |

*Note.* Unstandardized coefficients (*B*), standard errors (*SE*), *z*-values, *p*-values, 95% bootstrap confidence intervals, and standardized coefficients ( $\beta$ ) are reported. Indirect effects were estimated using 1,000 bootstrap samples.

For self-related learning, belief updating following negative prediction errors was reduced with increased narcissistic personality traits (direct negative effect on negative self-related learning rates; NARQ → Self α_PE-_: *B* = -0.080, *SE* = 0.040, *z* = -2.00, *p* = .045, 95% CI [-0.158, -0.002]; see Figure 2). This was not mediated by pride. When feedback was better than expected, narcissistic personality traits predicted stronger self-belief updating with greater pride (indirect-only mediation effect on positive self-related learning rates via pride; NARQ → Pride → Self α_PE+_: *B* = 0.028, *SE* = 0.011, *z* = 2.65, *p* = .008, 95% CI [0.009, 0.050]; direct effect NARQ → Self α_PE+_ n.s.: *B* = −0.027, *SE* = 0.020, *z* = -1.34, *p* = .181, 95% CI [-0.094, 0.001]).

For other-related learning, narcissistic personality traits were related to more negative belief updates about the other person with greater pride (indirect-only mediation on negative other-related learning rates; NARQ → Pride → Other α_PE-_: *B* = 0.024, *SE* = 0.012, *z* = 2.09, *p* = .037, 95% CI [0.003, 0.048]; direct effect NARQ → Other α_PE-_n.s.: *B* = 0.009, *SE* = 0.029, *z* = 0.31, *p* = .755, 95% CI [-0.044, 0.070]). Positive learning rates for the other person were neither related to narcissistic traits nor pride in this model (for a full representation of all direct and indirect effects see Table 2 and for a control analysis including group, see Supplementary Table 6).

Overall, the results suggest that pride plays a mediating role in how narcissistic personality traits are associated with how people form new beliefs about themselves and others. In addition to updating less negatively about oneself with stronger narcissistic traits, stronger pride experiences in people with higher narcissistic traits are linked to increased positive self-related learning. Both can be seen as ways of self-enhancement. Furthermore, a stronger pride experience mediates the link between higher narcissistic personality traits and more negative learning about the other player, which can be interpreted as a devaluation of others.

## Discussion

This study examined the relationship between narcissistic personality traits and both computational processes and affective mechanisms of belief formation. The present findings suggest self-serving biases in belief updating are associated with narcissistic personality traits: First, in line with the idea of self-enhancement and grandiosity, individuals higher in narcissistic personality traits showed generally elevated performance expectations for themselves relative to others, which further increased over time, consistent with a pattern of greater positive self-belief updating and reduced negative self-belief updating. Second, this motivational pattern was also reflected at the computational level: the self-related negativity bias in learning rates was reduced in individuals with higher narcissistic personality traits, and third, stronger narcissistic personality traits were directly associated with reduced updating of self-related beliefs following worse-than-expected feedback, as well as stronger updating following better-than-expected feedback, indirectly mediated by experiences of pride. Both effects can be viewed as ways to uphold one’s grandiose self-image: one bolsters against disconfirming, self-threatening information, and the other boosts positive self-views via experiences of pride. Fourth, narcissistic personality traits predicted stronger updating of beliefs about others following worse-than-expected feedback, again indirectly mediated by pride. This effect can be viewed as a higher tendency to devalue others with increasing narcissistic personality traits. Together, these findings point toward multiple, converging mechanisms through which self-belief formation is shaped by narcissistic personality traits, spanning baseline expectation biases, asymmetric sensitivity to prediction errors, and affective pathways, all with implications for one’s status and self-worth maintenance.

### Reduced integration of negative self-related information

The reduced updating following worse-than-expected self-related feedback aligns with conceptualizations of narcissism as involving defensive self-regulation and self-enhancement processes^30^. Usually, negative information has a stronger impact on self-perception than positive information^31^, as it is often perceived as more extreme, intense, and surprising^31–33^. In performance contexts, such as the present learning paradigm, numerous studies have consistently demonstrated a negativity bias, with stronger self-related updating after negative than positive feedback^6–8^, suggesting that negative information may be especially salient when performance feedback bears on the self. For individuals high in narcissistic traits, however, incorporating such information may pose a greater threat to a positive self-image. In the performance context of the present learning paradigm, the possibility of failure may have further heightened attention to personal mistakes^8^, increasing the relevance of self-protective processes. The observed pattern therefore suggests an interplay between contextual factors and personality traits in shaping feedback processing. Rather than reflecting insensitivity to feedback per se, reduced updating among individuals with higher narcissistic traits may reflect motivated resistance to information that threatens a positive self-belief, thereby supporting the maintenance of a grandiose self-image^1^.

### Pride as an affective gateway to self-enhancing learning

In contrast to the direct link between narcissistic traits and attenuated updating following negative feedback, the association between narcissistic traits and increased learning from positive feedback was mediated by pride. This pattern indicates that enhanced learning from better-than-expected information is systematically linked to affective experience rather than being purely cognitive. Pride has been conceptualized as a self-conscious emotion that is closely linked to achievement, competence, and social rank^34–36^. Prior evidence also specifically links grandiose narcissism to heightened pride^37,38^ and supports theoretical models that highlight pride as central to narcissistic self-regulation^27^. Apart from replicating the association between narcissism and heightened pride, the present findings also extend previous research demonstrating that affective states shape belief formation. Prior work has shown that the desire to maintain favorable beliefs, along with the emotions associated with them, can bias self-related learning^39^. Likewise, Müller-Pinzler et al. showed that stronger adjustments to negative feedback were accompanied by more embarrassment and less pride^7^. In the context of narcissism, pride may function as a reinforcement signal for positive self-related information, increasing the weight of better-than-expected feedback and strengthening belief updating when outcomes confirm one’s own competence.

However, the directionality of these processes remains open, as stronger integration of better-than-expected feedback may itself increase experienced pride. Reciprocal processes may also operate, whereby pride and belief updating dynamically reinforce each other. Thus, narcissism may be associated with stronger experiences of pride, which in turn coincide with enhanced integration of positive self-related information, potentially reflecting a mutually reinforcing relationship between affective responses and belief updating. It should be noted that mediation in non-experimental designs does not permit a causal inference, and that the temporal ordering of the measures places further constraints on causal interpretation: pride was assessed concurrently with the learning task, whereas narcissistic traits were assessed only afterward, although narcissistic traits are generally considered relatively stable personality characteristics.

### Updating beliefs about others in narcissistic personality traits: relative status dynamics

Narcissistic traits indirectly predicted stronger updating about others following worse-than-expected feedback in an indirect-only mediation via pride. This is particularly interesting as negative information about others may imply a relative status advantage. Narcissism is also closely tied to a status motive, whereby competition serves to demonstrate superiority and stabilize self-esteem^40,41^. In the formation of beliefs about others, self-enhancement associated with narcissistic traits is accompanied by the devaluation of others, particularly those who outperform the individual in ego-relevant domains. Such tendencies may, in turn, contribute to biased processing of feedback about others^42–44^.

These rivalry aspects of narcissism are consistently linked to psychological dysfunction, including depression and substance use disorders^45,46^, as well as adverse interpersonal outcomes, such as hostility and interpersonal conflict^47,48^, and reflect a more maladaptive facet^1^. The clinical part of the sample was recruited from a day clinic specializing in interpersonal problems in chronic depression. The tendency to devalue others may contribute to interpersonal difficulties and could represent one pathway through which narcissistic traits are linked to psychological dysfunction.

In the present study, the link between higher narcissism and more negative belief updating about others emerged only when mediated via pride. This might indicate that heightened pride in narcissism may not only predict more positive self-related belief updating but also a comparative evaluative shift in which the self is elevated relative to others. Because pride is often tied to perceptions of relative standing^49^, the experience of pride may not only reinforce positive self-related learning but also amplify contrast effects in social comparison, leading to more negative updating about others. Also here, the directionality remains open, with pride and belief updating potentially reinforcing each other. Future research could test whether a direct competitive context with feedback in comparison to the other participant, rather than a reference group (e.g., “You estimated 30% better than Tim”), is more effective in triggering the status motive and thus, a more negatively biased learning about the other person.

Notably, the present findings also highlight the value of implicit, cover story-based learning paradigms for investigating narcissistic self-regulation. The paradigm used in the present study is particularly well suited for examining narcissistic self-regulation processes because self-serving biases are not assessed explicitly through self-reports but are instead captured implicitly through trial-by-trial belief updating in response to manipulated performance feedback. This approach allows for the identification of self-serving biases without relying on participants’ conscious self-evaluations, thereby reducing potential distortions resulting from social desirability or strategic self-presentation, which may be especially relevant in the context of narcissism ^50^. Beyond preventing deliberate response biases, the present paradigm also captures processes that occur unconsciously. Such a methodological approach aligns closely with theoretical accounts of narcissism proposed by Morf and Rhodewalt, conceptualizing narcissistic self-regulation as a largely automatic and nonreflective process triggered by situational cues related to self-enhancement or threats to self-affirmation^25^. From this perspective, narcissism constitutes an affect-based disposition driven by the chronic need to maintain self-esteem, resulting in a self-system that remains vigilantly attuned to opportunities for self-enhancement. Consistent with this framework, pride emerged as a mediating factor in the present study, further underscoring the role of affective processes in narcissistic self-regulation. Accordingly, the present measures of belief updating may be particularly sensitive to the dynamic interplay of cognitive, affective, and self-evaluative processes underlying self-belief formation and self-maintenance in narcissism. As such, they may provide a valuable window into the cognitive and affective mechanisms through which narcissistic individuals maintain and reproduce favorable self-views.

## Conclusion

In conclusion, this study provides new insights into the role of narcissistic personality traits in self- and other-related belief formation. Unlike prior work, it focused on the development of a domain-specific self-concept to shed light on the dynamics of narcissism and forming beliefs about oneself and others. Taken together, the findings suggest a self-serving, status-sensitive, and affectively modulated shift in belief updating related to narcissistic traits. This pattern fits integrative models conceptualizing narcissism as a dynamic self-regulatory system organized around status pursuit and maintenance^51^. Importantly, the mediation findings indicate that the affective experience, specifically pride, plays a central role in modulating how feedback is incorporated into beliefs, extending theoretical models of narcissism that emphasize dynamic self-regulation^25^ by empirically testing how affective states shape belief updating.

## Methods

### Participants

The data are based on a sample partially published in Czekalla et al. (2026)^8^. It consists of a clinical sample of individuals diagnosed with depression, a healthy control group who performed the task in the MRI^8^, and an additional clinical sample that performed the task outside the MRI scanner. All participants were above the age of 18 and fluent in German. The clinical sample (n = 65, 26 females, aged 19-60, M = 37.4, SD = 11.5) included individuals diagnosed with depression (ICD-10^52^ codes: F32.1/2, F33.1/2). Participants were excluded from the clinical sample if they presented acute suicidality, an acute psychotic state, schizophrenia, schizotypal disorder or delusional disorder, bipolar disorder, primary diagnosis of substance abuse or substance dependence, marked personality disorder diagnoses in the narcissistic, histrionic, or borderline domains, or any neurological condition. The sample was recruited from patients receiving treatment at the Department of Psychiatry and Psychotherapy at the University Hospital Lübeck’s day hospital.

The healthy sample (n = 32, 10 females, aged 18-55, M = 34.3, SD = 10.2) was recruited via the University of Lübeck’s email list and flyers. Exclusion criteria were neurological or mental illnesses, previous participation in related studies, or advanced psychology students. Eligibility was verified in an interview via phone before the study. For this study, both samples were combined to maximize the sample size and make use of a wide variance in trait narcissism (for a detailed description of sample characteristics, see Supplementary Tables 7 and 8).

Originally, 116 participants were recruited in total. 19 had to be excluded due to technical issues, early study termination, or specific questionnaire incompleteness, resulting in a final sample of 97 participants. The study participants received an expense allowance of 10 euros per hour, i.e., a total of 25 euros for full participation in the measurement. Psychology students could also be compensated with course credit for participation. To motivate careful responding, participants were supposedly able to earn up to an additional 5 euros for accurate predictions of their estimation performance; this amount was fully paid out after the cover story was revealed.

The study was approved by the ethics committee of the University of Lübeck and conducted in accordance with the ethical guidelines of the American Psychological Association. All participants provided written informed consent before participating.

### General Procedure. Learning Of Own Performance task

The LOOP task^6–8^ is aimed at measuring the formation of self-beliefs about one’s abilities compared to forming beliefs about another person. More precisely, the focus is on identifying how positive and negative performance feedback are weighted to capture interindividual differences in belief-updating biases. The task was masked as a measure of cognitive estimation. Two subjects were invited; one completed the tasks in the MRI scanner, while the other performed the task as a behavioral study. In the case of no second participant available, a confederate was presented as a second participant. Before the task, participants gave written consent, completed questionnaires (not included in this paper), and received detailed instructions concerning the tasks.

The paradigm consisted of a total of 80 trials. Each participant was randomly assigned to two of the four estimation categories (e.g., heights of buildings, weights of animals), while the remaining two categories were completed by the other participant. Thus, in two estimation categories, the feedback referred to the participant’s own estimation performance (Self condition), whereas in the remaining two categories it referred to the other participant’s performance (Other condition). One of the two estimation categories was associated with predominantly positive feedback (High ability condition) for each agent condition, while the other estimation category was associated with negative feedback (Low ability condition). This design enabled participants to gradually learn that either they or the other person performed well in one estimation category but poorly in the other. It resulted in four feedback conditions (Agent [Self vs Other] x Ability [High vs Low]) with 20 trials each. The estimation categories were counterbalanced between the ability condition and the agent condition.

At the beginning of each trial, the participants saw a cue with the upcoming estimation category and whose turn it was (e.g., “You”, “Weights”). Afterward, the participant assessed how good they expected their own or the other person’s performance to be (performance expectation; EXP). The performance expectation was indicated as a percentile rank on a scale from zero to 100 in comparison to a supposed reference group of 350 test subjects that had been surveyed in advance. There was no time limit for this rating. To motivate careful rating of the performance expectation, participants were informed that they would be paid up to 6 cents for an accurate performance expectation rating. The participants were then presented with an image from one of the categories along with the corresponding estimation question. Within ten seconds, the participants were asked to estimate, e.g., how high the depicted building was. The given range of the dimensional answer scale was adjusted depending on the estimation question. After submitting an estimate, the participants received feedback. The feedback was displayed for three seconds and presented as a visual percentage rank in comparison to the reference group (e.g., “You are/ Riley is better than 55 % of the reference group”).

The feedback was independent of the actual estimation performance and manipulated according to the ability conditions as follows: The mean value from the performance expectations of the participants’ last five trials was calculated as a measure of the current ability belief. A preprogramed amount of deviation was added or subtracted to this mean value, which resulted in the feedback presented. For the high ability condition, feedback was better than expected in 70% of cases (i.e., positive prediction error) and worse than expected in 30% (i.e., negative prediction error), whereas in the low ability condition, this pattern was reversed. The aim was to balance the prediction error across the estimation categories. Furthermore, the participants’ current emotional state was recorded every 20 trials. The participants rated their perceived level of embarrassment, pride, joy, and arousal on a continuous scale from not at all (coded as zero) to very strongly (coded as 100). A follow-up survey then included the Narcissistic Admiration and Rivalry Questionnaire^1^ among other questionnaires (not included).

### Capturing the level of narcissistic traits

Trait narcissism was assessed using the German version of the Narcissistic Admiration and Rivalry Questionnaire (NARQ^1^). It captures grandiose narcissism as a two-dimensional construct, containing narcissistic need for admiration and narcissistic rivalry. To analyze the data in the current study, the two scales of admiration and rivalry were combined, and a total score for the expression of narcissism was determined for each participant. The mean value of the answers across all items was calculated for this purpose.

## Statistical analysis

### Model-agnostic analysis

First, we performed a model-agnostic analysis on the participants’ expected performance ratings for each trial to capture the basic effects of belief updating in our observed data. Performance expectation ratings were analyzed using a linear mixed model estimated with restricted maximum likelihood. Fixed effects comprised Ability (High/Low), Agent (Self/Other), NARQ, and Trial (1–20, treated as continuous). To account for individual differences, random intercepts and random slopes for all within-subject factors (Ability, Agent, and Trial) were specified per participant. Further, another linear mixed model was implemented that additionally included Group (MDD/ CON) as a factor (fixed effect) to control for potential differences between groups.

### Computational modeling

Dynamic changes in performance expectation ratings were modeled using PE delta-rule update equations (adapted Rescorla–Wagner model^53^). The model space and the procedure of model fitting and selection have been implemented and further developed in our previous studies^6,7^ and correspond to the procedure described in the previous study that includes a subgroup of the current sample^8^. The model space included three main approaches (see also Supplementary Table 3): the **Unity Model** (single learning rate), the **Valence Model** (distinct learning rates for positive vs. negative prediction errors), and the **Ability Model** (distinct learning rates for ability conditions). A **Mean Model** (M0) served as a baseline, assuming stable values. Building on the Valence Model (M6), we added an individual weighting factor to account for the reduced perceived likelihood of extreme feedback values (i.e., percentiles near 0% or 100%)^54^. Each feedback percentile was assigned a weight based on the relative probability density of a normal distribution, such that feedback near the mean received higher weights than extreme values. The strength of this downweighting was captured by an individually fitted parameter w. This weighted Valence Model (M7) was the winning model in our last study^7^. The normal decay (ND) weighted by the weighting factor w was introduced in the learning models in the following way:

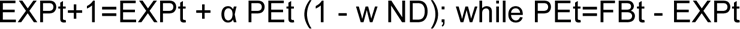

Learning rates were estimated either separately for Self vs. Other or across both agents. The initial beliefs about the own and the other participant’s performance (EXP1) were estimated as free parameters separately for Self and Other in both Ability conditions, resulting in four additional model parameters. All models in the model space were fitted and compared as described below.

Individual learning parameters were estimated via Markov Chain Monte Carlo sampling implemented in RStan. Three chains each contributed 2400 posterior samples, preceded by 1000 burn-in samples with a thinning factor of 3. Model convergence was evaluated using *R̂* statistics^55^; effective sample sizes (neff) > 1500 for the majority of parameters and subjects. For group-level analyses, individual parameter estimates were derived from the mean of the posterior distribution.

Model selection was based on predictive accuracy, assessed via leave-one-out cross-validation using Pareto smoothed importance sampling (PSIS-LOO)^56^. Reliability of the PSIS-LOO estimates was evaluated through *k̂* diagnostics; only a small number of trials yielded values indicative of potentially unreliable estimates (Supplementary Table 3). The winning model was validated by replicating the model-agnostic analyses with its predicted data (see Supplementary Note 1, Supplementary Table 9).

### Analyses of learning rates

Learning rates derived from the weighted Valence Model (winning computational model) were analyzed in a linear mixed model with a restricted maximum likelihood estimation. The model included the factors PE Valence (Pos/ Neg) x Agent (Self/Other) x NARQ. Intercept, PE Valence, Agent, and NARQ were modeled as fixed effects. Random intercepts and a random slope for Agent were included for each participant. A random slope for PE Valence was not included because the model failed to converge when both random slopes were specified. Further, another linear mixed model was implemented that additionally included Group (MDD/ CON) instead of NARQ as a factor (fixed effect) to control for and assess potential differences between groups.

### Path analytic analyses

To assess whether the association between narcissistic personality traits and altered learning about the own or others’ abilities was mediated by the experience of pride (average pride ratings during the task), we conducted mediation analyses using path-analytic models estimated in the R (version 4.5.2) lavaan 0.6.21 package^57^.

Specifically, narcissism (NARQ scores, global scale) was specified as a predictor of pride. Pride (mediator) was then modeled to predict the four learning rates derived from the winning computational model as outcome variables (learning rates for positive self- and other-related prediction errors [Self α_PE+_; Other α_PE+_] as well as negative self- and other-related prediction errors [Self α_PE-_; Other α_PE-_]). Direct effects of narcissism on each outcome were included to test for partial mediation. All outcome variables were estimated simultaneously, and their residuals were allowed to correlate. Indirect effects for each outcome were defined as the product of the corresponding paths from the predictor towards the mediator and from the mediator towards the outcome and were estimated within the model (see Figure 2). As the sample included a group of individuals with chronic depression (MDD) and a healthy control group (CON), a second model was set up to control for potential effects of group. The main path-analytic model described above was extended by including Group as an additional predictor alongside narcissism. Group was dummy-coded, and the two predictors were allowed to covary.

Standard errors and confidence intervals were obtained using bootstrap resampling (1,000 samples). Models were estimated using maximum likelihood with a mean structure included. As the specified mediation model was just-identified (df = 0), overall model fit indices were not informative and are therefore not reported.

## Data availability

The data used to generate the analyses are available at Zenodo (https://doi.org/10.5281/zenodo.23415012).

## Code availability

The analysis code is available on GitHub (https://github.com/NoCze/narcissismLOOP) and archived on Zenodo (https://doi.org/10.5281/zenodo.23415012). A pre-rendered overview of all results is provided as an R Quarto HTML output document within the repository. R Stan scripts to estimate the computational models are available from the corresponding author upon reasonable request.

## Acknowledgements

We are grateful to Rebecca Rocksien, Katharina Ohm, Ann-Kristin Schmidt, Jovana Lehmann-Grube, Alica Steinert, Leonie Christine Pasquier, and Yana Schwarze for their help with data collection.

## Author contributions

LM-P, SK, FMP, NC conceived and designed the experiments. NC and AS collected the data, with the consultation of JPK to recruit the clinical sample. MM, NC, LM-P, and LR analyzed the data. MM, AVM, AS, DSS, FMP, SK, LM-P, and NC discussed the data analyses and interpretation of the results. MM wrote the original draft of the manuscript. NC, LM-P, and SK supervised and edited the paper. AVM, AS, JPK, IW, DSS, and FMP reviewed and edited the paper.

