## Supplementary_material for "Self-enhancement and devaluation of others during belief formation are linked to narcissistic traits"

Matthiesen et al.

### Supplementary Notes

#### Supplementary Note 1: Posterior predictive checks - Behavioral analyses on the predicted data.

We repeated the same analyses as for the behavioral data (linear mixed model) using the predicted data from the winning model to see whether our winning model captured the core effects in our model-agnostic analysis. We could reproduce the main effects from the model-agnostic analysis with the predicted data (main effect Ability condition: $t(114.8)=-2.35$, $p=.020$; main effect Agent condition: $t(109.4)=-3.50$, $p=.001$; interaction Trial x Ability condition: $t(7368)=-37.97$, $p<.001$; interaction Trial x Agent condition $t(7368)=-9.91$, $p<.001$). The effects related to narcissistic personality traits could also be reproduced (Agent × NARQ interaction $t(111)=2.56$, $p=.012$, Agent × Trial × NARQ interaction $t(7364.0)=4.37$, $p<.001$). This confirms that the winning model recapitulates the main effects in our data (see Supplementary Table 9 for all results).

#### Supplementary Note 2: Path-analytic model controlling for potential differences between groups

As the sample included a group of individuals with chronic depression and a healthy control group, another path-analytic model was set up to control for potential effects of group (MDD/ CON). This model mostly replicated the main findings. There was a significant indirect effect of narcissism via pride for self-related positive learning rates and a direct effect of narcissism on self-related negative learning rates. The indirect effect of narcissism on other-related negative learning rates via pride showed a trend-wise effect when controlling for Group (NARQ → Pride → Other $\alpha_{PE-}$: $B = 0.021$, $SE = 0.011$, $z = 1.94$, $p = .052$, 95% CI [0.001, 0.044]). There were no significant effects of Group (see Supplementary Table 6 for a full representation of all direct and indirect effects).

### Supplementary Tables

*Supplementary Table 1.* Model-agnostic analysis of belief-updating behavior

| | *Estimate* | *SE* | 95% CI | | *df* | *t* | *p* |
|---|---|---|---|---|---|---|---|
| | | | lower | upper | | | |
| (Intercept) | 55.88 | 0.98 | 53.97 | 57.80 | 114.4 | 57.13 | < .001 |
| Trial | 0.49 | 0.04 | 0.40 | 0.57 | 252.4 | 11.46 | < .001 |
| Agent [Self] | -4.81 | 1.24 | -7.25 | -2.38 | 131.8 | -3.87 | < .001 |
| Ability [Low] | -4.13 | 1.26 | -6.61 | -1.66 | 130.9 | -3.27 | .001 |
| NARQ | 2.50 | 1.84 | -1.10 | 6.10 | 114.4 | 1.36 | .176 |
| Agent x Trial | -0.22 | 0.04 | -0.31 | -0.14 | 7363.9 | -5.18 | < .001 |
| Ability x Trial | -0.90 | 0.04 | -0.98 | -0.81 | 7363.9 | -20.81 | < .001 |
| Ability x Agent | -0.03 | 0.73 | -1.46 | 1.39 | 7363.9 | -0.05 | .962 |
| Trial x NARQ | -0.16 | 0.08 | -0.31 | 0.00 | 252.4 | -1.96 | .052 |
| Agent x NARQ | 5.24 | 2.33 | 0.67 | 9.81 | 131.8 | 2.25 | .026 |
| Ability x NARQ | -3.12 | 2.37 | -7.77 | 1.52 | 130.9 | -1.32 | .190 |
| Ability x Agent x Trial | -0.15 | 0.06 | -0.27 | -0.03 | 7363.9 | -2.52 | .012 |
| Agent x Trial x NARQ | 0.26 | 0.08 | 0.11 | 0.42 | 7363.9 | 3.26 | .001 |
| Ability x Trial x NARQ | -0.03 | 0.08 | -0.19 | 0.12 | 7363.9 | -0.43 | .665 |
| Ability x Agent x NARQ | 2.71 | 1.37 | 0.03 | 5.39 | 7363.9 | 1.98 | .048 |
| Ability x Agent x Trial x NARQ | -0.06 | 0.11 | -0.28 | 0.16 | 7363.9 | -0.53 | .598 |

*Note.* Trial (continuous, 1-20) x Agent [Self, Other] x Ability condition [High, Low] x NARQ linear mixed-effects model fit by maximum likelihood. Dependent variable: trial-by-trial ratings of expected performance. Beta-estimate, *SE* = standard error, *CI* = confidence interval, *df* = degrees of freedom, *t* = t-value.

*Supplementary Table 2.* Model-agnostic analysis of belief-updating behavior including Group

| | *Estimate* | *SE* | 95% CI | | *df* | *t* | *p* |
|---|---|---|---|---|---|---|---|
| | | | lower | upper | | | |
| (Intercept) | 57.43 | 1.69 | 54.12 | 60.75 | 114.5 | 33.93 | <.001 |
| Trial | 0.52 | 0.07 | 0.38 | 0.67 | 250.6 | 7.07 | <.001 |
| Agent [Self] | -4.42 | 2.32 | -8.97 | 0.12 | 126.1 | -1.91 | .059 |
| Ability [Low] | -5.49 | 2.19 | -9.79 | -1.20 | 131.1 | -2.51 | .013 |
| Group [MDD] | -2.32 | 2.07 | -6.37 | 1.74 | 114.5 | -1.12 | .265 |
| Agent x Trial | -0.20 | 0.07 | -0.35 | -0.06 | 7364.1 | -2.70 | .007 |
| Ability x Trial | -1.08 | 0.07 | -1.23 | -0.93 | 7364.1 | -14.39 | <.001 |
| Ability x Agent | 0.81 | 1.27 | -1.68 | 3.30 | 7364.1 | 0.64 | .525 |
| Trial x Group | -0.06 | 0.09 | -0.23 | 0.12 | 250.6 | -0.62 | .538 |
| Agent x Group | -0.58 | 2.83 | -6.13 | 4.97 | 126.1 | -0.20 | .838 |
| Ability x Group | 2.03 | 2.68 | -3.21 | 7.28 | 131.1 | 0.76 | .449 |
| Ability x Agent x Trial | -0.07 | 0.11 | -0.28 | 0.14 | 7364.1 | -0.64 | .525 |
| Agent x Trial x Group | -0.03 | 0.09 | -0.21 | 0.15 | 7364.1 | -0.33 | .740 |
| Ability x Trial x Group | 0.27 | 0.09 | 0.09 | 0.45 | 7364.1 | 2.98 | .003 |
| Ability x Agent x Group | -1.26 | 1.55 | -4.30 | 1.78 | 7364.1 | -0.81 | .418 |
| Ability x Agent x Trial x Group | -0.13 | 0.13 | -0.38 | 0.13 | 7364.1 | -0.99 | .320 |

*Note.* Trial (continuous, 1-20) x Agent [Self, Other] x Ability condition [High, Low] x Group [MDD, CON] linear mixed-effects model fit by restricted maximum likelihood. Dependent variable: trial-by-trial ratings of expected performance. Beta-estimate, *SE* = standard error, *CI* = confidence interval, *df* = degrees of freedom, *t* = t-value.

*Supplementary Table 3.* Model comparison

| *Model* | *PSIS-LOO* | *LOO-SE* | *LOO-Diff (SE-Diff)* | *% of $\hat{k}$> 0.7* | *Num. param.* |
|---|---|---|---|---|---|
| Mean Model (M0) | -4140.3 | 343.1 | -1668.9 (208.1) | 0.07 | 4 |
| Self = Other | | | | | |
| Unity Model (M1) | -3370.5 | 407.6 | -899.1 (157.8) | 1.24 | 5 |
| Ability Model (M2) | -3207.7 | 414.3 | -736.3 (142.9) | 1.45 | 6 |
| Valence Model (M3) | -3109.6 | 427.5 | -638.3 (109.9) | 0.89 | 6 |
| Self ≠ Other | | | | | |
| Unity Model (M4) | -3241.6 | 412.4 | -770.3 (124.3) | 1.10 | 6 |
| Ability Model (M5) | -3026.6 | 414.4 | -555.2 (107.0) | 2.57 | 8 |
| Valence Model (M6) | -2639.0 | 441.1 | -167.7 (36.7) | 0.84 | 9 |
| Weighted Valence Model (M7) | -2471.3 | 439.9 | - | 2.15 | 8 |

Note. PSIS-LOO = sum PSIS-LOO, approximate leave-one-out cross-validation (LOO) using Pareto-smoothed importance sampling (PSIS); LOO-SE = Standard error of PSIS-LOO; LOO-Diff (SE-Diff) = Difference in expected predictive accuracy (PSIS-LOO) for all models from the model with the highest PSIS-LOO (weighted Valence Model) and standard errors of differences; percentage of $\hat{k}$ - estimated shape parameters of the generalized Pareto distribution - exceeding 0.7 (all according to Vehtari et al., 2016); Num. param. = number of estimated parameters in the model; Self = Other: same learning rates for Self and Other, Self ≠ Other: separate learning rates for Self and Other.

*Supplementary Table 4.* Analysis of learning rates

| | *Estimate* | *SE* | 95% CI | | *df* | *t* | *p* |
|---|---|---|---|---|---|---|---|
| | | | lower | upper | | | |
| (Intercept) | 0.20 | 0.016 | 0.17 | 0.23 | 163.1 | 12.30 | < .001 |
| PE Valence [Pos] | 0.04 | 0.017 | 0.01 | 0.07 | 190.0 | 2.50 | .013 |
| Agent [Self] | 0.11 | 0.019 | 0.08 | 0.15 | 211.8 | 5.96 | < .001 |
| NARQ | 0.03 | 0.031 | -0.03 | 0.09 | 163.1 | 1.08 | .282 |
| PE Valence x Agent | -0.17 | 0.024 | -0.21 | -0.12 | 190.0 | -6.98 | < .001 |
| PE Valence x NARQ | -0.03 | 0.031 | -0.09 | 0.03 | 190.0 | -1.03 | .307 |
| Agent x NARQ | -0.15 | 0.036 | -0.22 | -0.08 | 211.8 | -4.19 | < .001 |
| PE Valence x Agent x NARQ | 0.15 | 0.045 | 0.06 | 0.24 | 190.0 | 3.36 | < .001 |

*Note.* PE Valence [Pos, Neg] x Agent [Self, Other] x NARQ linear mixed-effects model fit by restricted maximum likelihood. Dependent variable: learning rates. Beta-estimate, *SE* = standard error, *CI* = confidence interval, *df* = degrees of freedom, *t* = t-value.

*Supplementary Table 5.* Analysis of learning rates including Group

| | *Estimate* | *SE* | 95% CI | | *df* | *t* | *p* |
|---|---|---|---|---|---|---|---|
| | | | lower | upper | | | |
| (Intercept) | 0.22 | 0.029 | 0.17 | 0.28 | 168.4 | 7.77 | <.001 |
| PE Valence [Pos] | 0.04 | 0.030 | -0.02 | 0.10 | 190.0 | 1.30 | .196 |
| Agent [Self] | 0.07 | 0.034 | 0.01 | 0.14 | 213.5 | 2.11 | .036 |
| Group [MDD] | -0.03 | 0.035 | -0.10 | 0.04 | 168.4 | -0.92 | .359 |
| PE Valence x Agent | -0.14 | 0.043 | -0.23 | -0.06 | 190.0 | -3.31 | .001 |
| PE Valence x Group | 0.00 | 0.037 | -0.07 | 0.08 | 190.0 | 0.11 | .916 |
| Agent x Group | 0.06 | 0.042 | -0.02 | 0.14 | 213.5 | 1.49 | .139 |
| PE Valence x Agent x Group | -0.04 | 0.052 | -0.14 | 0.07 | 190.0 | -0.68 | .500 |

*Note.* PE Valence [Pos, Neg] x Agent [Self, Other] x Group [MDD, CON] linear mixed-effects model fit by restricted maximum likelihood. Dependent variable: learning rates. Beta-estimate, *SE* = standard error, *CI* = confidence interval, *df* = degrees of freedom, *t* = t-value.

*Supplementary Table 6.* Path-analytic model including NARQ and Group

| | *B* | *SE* | 95% CI | | *z* | *p* | *β* |
|---|---|---|---|---|---|---|---|
| | | | lower | upper | | | |
| **Direct effects** | | | | | | | |
| NARQ → Pride | 0.768 | 0.173 | 0.433 | 1.116 | 4.43 | < .001 | 0.41 |
| Group → Pride | −0.301 | 0.196 | −0.689 | 0.083 | −1.54 | 0.124 | −0.14 |
| Pride → $\alpha_{PE+}$ Self | 0.034 | 0.012 | 0.012 | 0.058 | 2.93 | 0.003 | 0.33 |
| NARQ → $\alpha_{PE+}$ Self | −0.026 | 0.020 | −0.063 | 0.016 | −1.27 | 0.205 | −0.13 |
| Group → $\alpha_{PE+}$ Self | 0.009 | 0.021 | −0.032 | 0.051 | 0.40 | 0.688 | 0.04 |
| Pride → $\alpha_{PE-}$ Self | −0.045 | 0.026 | −0.099 | 0.002 | −1.73 | 0.084 | −0.23 |
| NARQ → $\alpha_{PE-}$ Self | −0.082 | 0.040 | −0.164 | −0.005 | −2.04 | 0.041 | −0.22 |
| Group → $\alpha_{PE-}$ Self | −0.015 | 0.037 | −0.087 | 0.060 | −0.39 | 0.697 | −0.03 |
| Pride → $\alpha_{PE+}$ Other | 0.022 | 0.017 | −0.010 | 0.056 | 1.28 | 0.200 | 0.14 |
| NARQ → $\alpha_{PE+}$ Other | −0.022 | 0.031 | −0.077 | 0.045 | −0.69 | 0.492 | −0.07 |
| Group → $\alpha_{PE+}$ Other | −0.023 | 0.033 | −0.088 | 0.042 | −0.71 | 0.478 | −0.07 |
| Pride → $\alpha_{PE-}$ Other | 0.028 | 0.014 | 0.001 | 0.056 | 1.97 | 0.048 | 0.20 |
| NARQ → $\alpha_{PE-}$ Other | 0.007 | 0.030 | −0.047 | 0.070 | 0.23 | 0.821 | 0.03 |
| Group → $\alpha_{PE-}$ Other | −0.017 | 0.032 | −0.079 | 0.045 | −0.53 | 0.598 | −0.06 |
| **Indirect effects** | | | | | | | |
| NARQ → Pride → $\alpha_{PE+}$ Self | 0.026 | 0.010 | 0.009 | 0.049 | 2.56 | 0.011 | 0.13 |
| NARQ → Pride → $\alpha_{PE-}$ Self | −0.035 | 0.022 | −0.083 | 0.002 | −1.56 | 0.118 | −0.09 |
| NARQ → Pride → $\alpha_{PE+}$ Other | 0.017 | 0.014 | −0.008 | 0.045 | 1.22 | 0.223 | 0.06 |
| NARQ → Pride → $\alpha_{PE-}$ Other | 0.021 | 0.011 | 0.001 | 0.044 | 1.94 | 0.052 | 0.08 |
| Group → Pride → $\alpha_{PE+}$ Self | −0.010 | 0.008 | −0.028 | 0.003 | −1.33 | 0.183 | −0.05 |
| Group → Pride → $\alpha_{PE-}$ Self | 0.014 | 0.014 | −0.003 | 0.049 | 0.96 | 0.338 | 0.03 |
| Group → Pride → $\alpha_{PE+}$ Other | −0.006 | 0.008 | −0.026 | 0.004 | −0.83 | 0.407 | −0.02 |
| Group → Pride → $\alpha_{PE-}$ Other | −0.008 | 0.008 | −0.029 | 0.002 | −1.04 | 0.299 | −0.03 |

*Note.* Unstandardized coefficients (*B*), standard errors (*SE*), *z*-values, *p*-values, 95% bootstrap confidence intervals, and standardized coefficients (β) are reported. Indirect effects were estimated using 1,000 bootstrap samples.

*Supplementary Table 7.* Sample Description

| | Total | CON | MDD |
|---|---|---|---|
| Sample Size | 97 | 32 | 65 |
| Gender (m, f) | 61/ 36 | 22/ 10 | 39/26 |
| Age, M (SD) | 36.4 (11.2) | 34.3 (10.2) | 37.4 (11.5) |
| Min/Max | 18.1/ 60.3 | 18.1/ 55.2 | 19.7/ 60.3 |
| NARQ, M (SD) | 1.88 (0.54) | 2.06 (0.52) | 1.80 (0.53) |
| Min/Max | 1.00/ 3.50 | 1.22/ 3.50 | 1.00/ 3.11 |
| BDI-V, M (SD) | 41.2 (23.4) | 13.9 (8.00) | 54.6 (15.3) |
| Min/ Max | 2/ 88 | 2/ 33 | 7/ 88 |

*Note.* CON = healthy control group, MDD = group of individuals diagnosed with major depressive disorder, M = mean value, SD = standard deviation, Min/Max = minimum/maximum, NARQ = Narcissistic Admiration and Rivalry Questionnaire (Back et al., 2013), BDI-V = simplified Becks Depression Inventory, sum scores (Schmitt et al., 2003)

*Supplementary Table 8.* F-diagnoses in clinical sample

| ICD-10 Code | Diagnosis | *n* |
|---|---|---|
| F33.2 | Recurrent depressive disorder, current episode severe without psychotic symptoms | 49 |
| F34.1 | Dysthymia | 24 |
| F33.1 | Recurrent depressive disorder, current episode moderate | 10 |
| F40.1 | Social phobias | 10 |
| F43.1 | Post-traumatic stress disorder | 7 |
| F60.6 | Anxious (avoidant) personality disorder | 4 |
| F32.2 | Severe depressive episode without psychotic symptoms | 3 |
| F41.1 | Generalized anxiety disorder | 3 |
| F60.5 | Obsessive-compulsive personality disorder | 3 |
| F41.0 | Panic disorder | 2 |
| F42.0 | Obsessive-compulsive disorder, predominantly obsessional thoughts | 2 |
| F10.1 | Mental and behavioral disorders due to alcohol, harmful use | 1 |
| F10.2 | Mental and behavioral disorders due to alcohol, dependence syndrome | 1 |
| F10.20 | Mental and behavioral disorders due to alcohol, dependence syndrome, currently abstinent | 1 |
| F12.1 | Mental and behavioral disorders due to cannabinoids, harmful use | 1 |
| F31.4 | Bipolar affective disorder, current episode severe depression without psychotic symptoms | 1 |
| F40.00 | Agoraphobia without panic disorder | 1 |
| F40.01 | Agoraphobia with panic disorder | 1 |
| F42.2 | Obsessive-compulsive disorder, mixed obsessional thoughts and acts | 1 |
| F42.8 | Other obsessive-compulsive disorders | 1 |
| F45.0 | Somatization disorder | 1 |
| F50.8 | Other eating disorders | 1 |
| F60.31 | Emotionally unstable personality disorder, borderline type | 1 |
| F61.0 | Mixed and other personality disorders | 1 |
| F90.0 | Attention-deficit hyperactivity disorder, predominantly inattentive type | 1 |

*Note.* Discharge diagnoses; comorbidities included; therefore, *n* counts may exceed the clinical sample size of *n*=65.

*Supplementary Table 9.* Analysis of model-predicted performance expectations

| | *Estimate* | *SE* | 95% CI | | *df* | *t* | *p* |
|---|---|---|---|---|---|---|---|
| | | | lower | upper | | | |
| (Intercept) | 55.18 | 0.94 | 53.35 | 57.01 | 103.5 | 59.00 | < .001 |
| Trial | 0.58 | 0.04 | 0.51 | 0.65 | 160.9 | 15.89 | < .001 |
| Agent [Self] | -4.48 | 1.19 | -6.81 | -2.15 | 110.5 | -3.77 | < .001 |
| Ability [Low] | -2.60 | 1.10 | -4.76 | -0.44 | 113.5 | -2.36 | .020 |
| NARQ | 2.09 | 1.76 | -1.35 | 5.53 | 103.5 | 1.19 | .236 |
| Agent x Trial | -0.28 | 0.03 | -0.34 | -0.23 | 7364.0 | -9.95 | < .001 |
| Ability x Trial | -1.09 | 0.03 | -1.14 | -1.03 | 7364.0 | -38.10 | < .001 |
| Ability x Agent | -0.99 | 0.48 | -1.93 | -0.04 | 7364.0 | -2.04 | .042 |
| Trial x NARQ | -0.13 | 0.07 | -0.27 | 0.00 | 160.9 | -1.95 | .053 |
| Agent x NARQ | 5.71 | 2.23 | 1.34 | 10.08 | 110.5 | 2.56 | .012 |
| Ability x NARQ | -2.66 | 2.07 | -6.71 | 1.40 | 113.5 | -1.28 | .202 |
| Ability x Agent x Trial | -0.02 | 0.04 | -0.10 | 0.06 | 7364.0 | -0.43 | .664 |
| Agent x Trial x NARQ | 0.23 | 0.05 | 0.13 | 0.34 | 7364.0 | 4.37 | < .001 |
| Ability x Trial x NARQ | -0.03 | 0.05 | -0.14 | 0.07 | 7364.0 | -0.65 | .516 |
| Ability x Agent x NARQ | 1.92 | 0.91 | 0.14 | 3.70 | 7364.0 | 2.11 | .035 |
| Ability x Agent x Trial x NARQ | -0.01 | 0.08 | -0.16 | 0.13 | 7364.0 | -0.18 | .856 |

*Note.* Trial (continuous, 1-20) x Agent [Self, Other] x Ability condition [High, Low] x NARQ linear mixed-effects model fit by restricted maximum likelihood. Dependent variable: predicted trial-by-trial values by the winning computational model. Beta-estimate, *SE* = standard error, *CI* = confidence interval, *df* = degrees of freedom, *t* = t-value.
